# Regulation of the lncRNA *malat1*/Egr1 Axis by Wnt, Notch and TGF-β signaling: A Key Mechanism in Retina Regeneration

**DOI:** 10.1101/2025.01.03.631187

**Authors:** Sharanya Premraj, Poonam Sharma, Mansi Chaudhary, Pooja Shukla, Kshitiz Yadav, Omkar Mahadeo Desai, Rajesh Ramachandran

## Abstract

Adult zebrafish retinas rely on the resident Müller glia to maintain homeostasis and enable regeneration. Retina regeneration remains incomplete in mammals despite extensive efforts to emulate zebrafish regenerative conditions. Many studies have examined the reprogramming of zebrafish Müller glia cells, which is necessary for regeneration driven by regeneration-associated gene expression. Here, we show that the lncRNA *malat1*, crucial for many biological functions, plays essential roles during retina regeneration. We demonstrate that *malat1* functions through an Egr-dependent axis, modulated by Wnt, Notch, and TGF-β signaling pathways, and is necessary for effective retina regeneration. Moreover, we uncover that the antisense lncRNA *talam1*, which regulates *malat1* availability, is differentially regulated in zebrafish and mice, highlighting species-specific gene regulatory mechanisms after retinal injury. Cells with active TGF-β signaling stabilize Malat1 in mice while the same signaling destabilizes *malat1* in zebrafish. Taken together, our work uncovers a new role for the *malat1*/Egr1 axis in necessitating retina regeneration, which may have important implications for differential regenerative ability in vertebrates.

## Introduction

Müller glia-dependent retina regeneration has been extensively studied in zebrafish and mice models (Jui & Goldman, 2024). Unlike mammals, vertebrates such as fishes and frogs have remarkable abilities to regenerate the retina (Sharma & Ramachandran, 2022). Retina regeneration adopts several gene regulatory networks associated with embryonic development, cancer, and wound healing (Goldman & Poss, 2020; Wong & Whited, 2020). Several regeneration-associated pathways that activate different gene regulatory networks essential to zebrafish retina regeneration are either suboptimal or antagonistic in the injured mice retina. Forced emulated zebrafish-specific retinal gene expression in mice often resulted in better regenerative response, suggesting the inevitable relevance of these regeneration-associated genes (Guo *et al*, 2023).

Although non-coding RNAs involve several biological phenomena (Dey *et al*, 2014; Hu *et al*, 2012), this study mainly focuses on the long non-coding RNA *malat1* (*metastasis-associated lung adenocarcinoma transcript 1*). It has been extensively demonstrated to be relevant to many biological functions, including embryonic development (Wu *et al*, 2019a), cellular proliferation (Shih *et al*, 2021), tissue differentiation (Yi *et al*, 2019), homeostasis (Li *et al*, 2021) , and cancer (Xu *et al*, 2024). Their roles during tissue regeneration have also been documented (Chen *et al*, 2017), but they remain unexplored in the context of retina regeneration.

Here, we investigated the regulation of *malat1* during zebrafish retina regeneration and its broader role in coordinating gene expression programs essential to mounting a robust regenerative response. During retina regeneration, we explored the link between *malat1* and different signaling pathways, such as Tgf-β, Wnt, and Delta-Notch signaling. Furthermore, we demonstrated that zebrafish and mice retinas exhibit distinct responses to injury, potentially driven by the Tgf-β signaling-mediated regulation of *malat1* in the injured retina of both species.

## Results

### The lncRNA *malat1* is differentially expressed in the regenerating retina

We explored the transcript levels of lncRNA *malat1* at different time points post-retinal injury by quantitative real-time PCR (qPCR) (Fig 1A). The *malat1* showed an upregulation in the reprogramming phase (15 hours post-injury (hpi)), and late differentiation phase (8 days post-injury (dpi)), compared to its levels in the uninjured retina. The overall levels of *malat1* are restored to uninjured levels at around 15 dpi. This expression pattern is verified by the RNA *in situ* hybridization done on retinal cross-sections at various times post-injury (Fig 1B). At 4 dpi, coinciding with the peak proliferation of retinal progenitors, *malat1* levels were comparable to those of the uninjured retina. A closer look at the RNA *in situ* hybridization at 4dpi reveals that *malat1* is absent in the proliferating BrdU^+^ retinal progenitors while expressing in the immediate neighboring cells at 4dpi and 7dpi (Fig 1C and Fig EV1A). Analysis of the proportion of proliferative *malat1*^+^ cells, indicated by proliferating cell nuclear antigen (PCNA) expression, reveals that both 4dpi and 7dpi harbor the *malat1* adjacent to the PCNA^+^ cells (Fig 1D). However, by 7dpi, some proliferated cells start expressing *malat1*.

**Figure 1.**
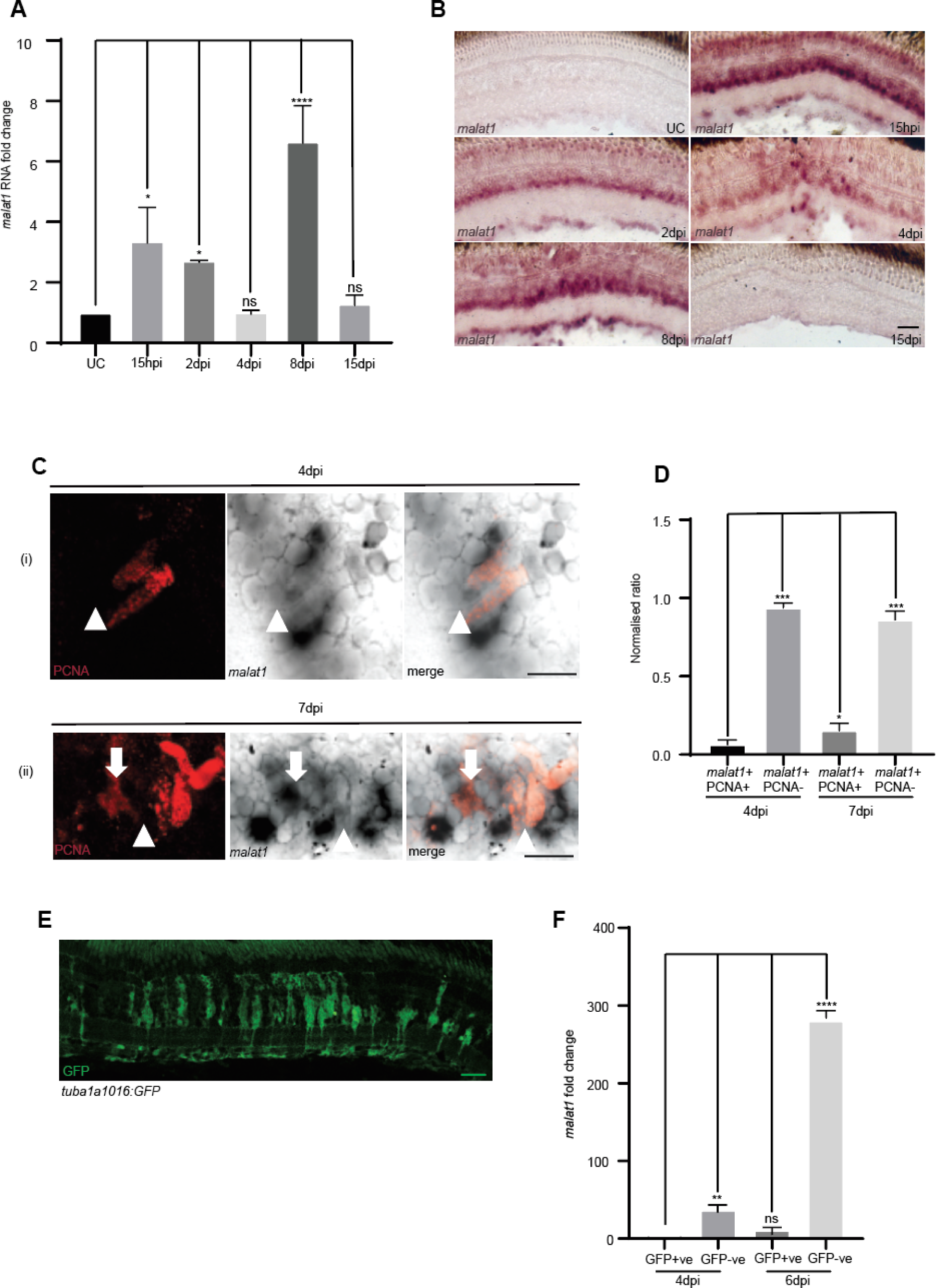
Temporal and spatial expression pattern of *malat1* during zebrafish retina regeneration. **(A)** qPCR analysis of *malat1* expression in the injured adult zebrafish retina at various time points post-injury. Data is presented as mean values ± standard deviation (SD), *p < 0.0003; n = 6 biological replicates. **(B)** 20X bright-field (BF) images of retinal cross-sections showing *malat1* localization after *in situ* hybridization using a probe specific for *malat1* in uninjured control (UC), 15 hours post-injury (hpi), 2 days post-injury (2 dpi), 4 dpi, 8 dpi, and 15 dpi retinae. **(C)** 60X bright-field image of a retinal cross-section at 4 dpi, showing *malat1* expression excluded from proliferating cells marked with PCNA (red). At 7 dpi, some inclusion is observed and quantified in **(D)**; *p < 0.005; n = 6 biological replicates. Arrows indicate cells with *malat1* inclusion, while arrowheads indicate *malat1* exclusion. **(E)** A representative image of a retinal cross-section from the *tuba1016:gfp* transgenic line harvested at 4 dpi, with GFP fluorescence highlighting actively proliferating cells. **(F)** qPCR analysis of *malat1* enrichment in GFP +ve (actively proliferating) and GFP-ve (neighboring) cells shows higher expression in neighboring cells compared to proliferating cells at 4 dpi and 6 dpi. Data is shown as mean values ± SD, *p < 0.05; n = 6 biological replicates; ns indicates non-significant. The scale bar represents 10 µm in panels **B**, **C**, and **E**.

To further validate this observation, a *1016tuba1a: GFP* transgenic line, expressing GFP in proliferating retinal progenitors, was used to sort cells and assess *malat1* expression levels in GFP+ proliferating cells as compared to GFP-cells (Fig 1E). qPCR analysis of *malat1* in GFP+ and GFP− cell populations revealed that *malat1* is predominantly expressed in GFP− cells (Fig 1F), confirming RNA *in situ* hybridization findings. Knockdown of *malat1* using antisense morpholinos (MO) in early developmental stages causes developmental defects and high mortality rates, highlighting this lncRNA’s inevitable roles (Fig EV1B and EV1C).

### Knockdown of *malat1* adversely affects the retinal progenitor proliferation

The *malat1*-targeting MO, with a tested efficacy (Fig EV2A), which adversely affected embryonic survival, was used to electroporate the injured zebrafish retina (Fig 2A). The number of retinal progenitors declined up to 50% upon *malat1* knockdown in a concentration-dependent manner in the 4dpi retina (Fig 2B and 2C). Since *malat1* expression is high at 15 hours post-injury (hpi) and 2dpi, a delayed knockdown strategy was implemented from 2 days post-injury (dpi) onwards. This delayed knockdown produced a similar reduction in retinal progenitors (Figs. 2D–2F). We performed a late *malat1* knockdown from 4 dpi to 8 dpi (the post-proliferative and differentiating phase) after marking proliferating cells with EdU at 4 dpi and BrdU at 8dpi (Fig 2G). The dual labeling was aimed at the following: (i) follow-up of the early progenitors after the *malat1* knockdown (EdU+), (ii) demarcating the early (EdU+) and late progenitors (BrdU+) after the *malat1* knockdown. Compared to the control, we saw a decline in the number of EdU^+^ and BrdU^+^ cells at 8dpi because of *malat1* knockdown (Fig 2H). These results suggest that *malat1* contributes to the induction and perpetual proliferation of retinal proliferation. We then explored the effect of overexpression of *malat1* in the regenerating retina. Overexpression of *malat1* increased proliferating retinal progenitors, hinting at its pro-proliferative effect. This effect was notably reduced when *malat1* was knocked down (Fig EV2B and EV2C). These observations show that *malat1* is essential for proper and adequate induction of retinal progenitors in the regenerating retina.

**Figure 2.**
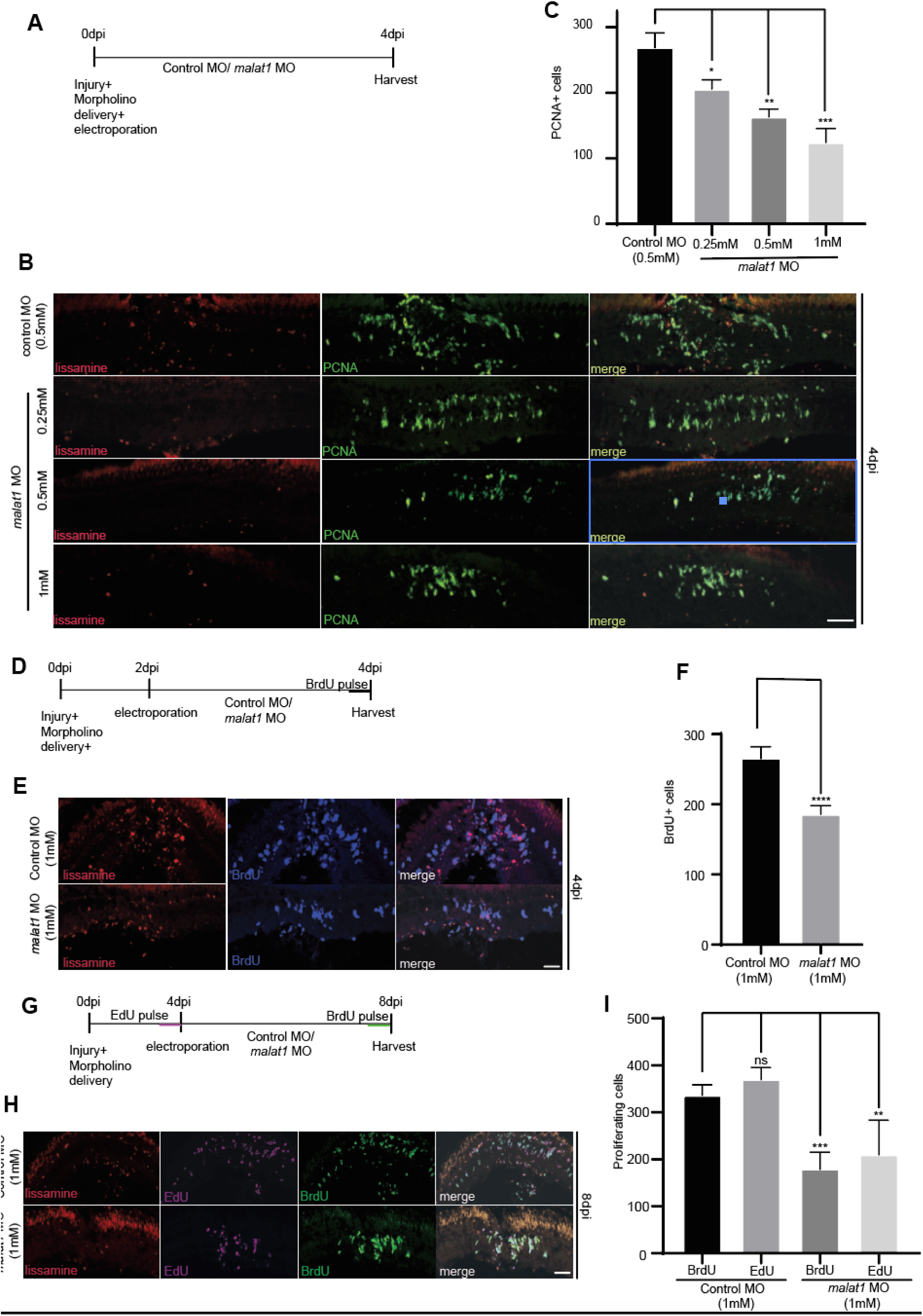
Impact of *malat1* knockdown on Müller Glia derived progenitor cell (MGPC) proliferation during different phases of zebrafish retina regeneration. *malat1* knockdown during proliferative, pre-proliferative, and late phases of retina regeneration results in reduced Müller glia-derived progenitor cell (MGPC) proliferation. A timeline schematic shows the injury sequence, electroporation, and BrdU/EdU pulsing (4 hours each), followed by tissue harvesting **(A, D, G). (B)** Retina injured and electroporated with lissamine-tagged morpholino (MO, red) displayed a concentration-dependent decrease in PCNA+ (green) cells when *malat1* was knocked down from 0 to 4 dpi, compared to control. Quantification is shown in **(C)**; *p < 0.001; n = 6 biological replicates. **(E)** Knockdown of *malat1* from 2 to 4 dpi led to a reduction in BrdU+ (cyan) cells, quantified in **(F)**; *p < 0.0005; n = 6 biological replicates. **(H)** Inhibition of *malat1* during the late proliferative phase (4 to 8 dpi) resulted in decreased BrdU+ (green) and EdU+ (pink) cells, with quantification in **(I)**; *p < 0.001; n = 6 biological replicates. ns indicates non-significant. Data is shown as mean values ± SD. The scale bar represents 10 µm **(B, E, H)**.

### *malat1* influences the expression of regeneration-associated genes

We saw the nuclear association of *malat1* revealed by fluorescence *in situ* hybridizations (FISH) with retinal cross-sections marked by DAPI, which revealed punctate nuclear expression of *malat1* (Fig 3A). Given its punctate pattern, we hypothesized that it may have a *trans* effect and investigated its association with epigenetic modifiers, including Ezh2, the PRC2 catalytic component, and Hdac1. We performed RNA immunoprecipitation (RIP) of 2dpi retinal extracts using anti-Ezh2 and anti-Hdac1 antibodies and probed for *malat1*. The results showed that *malat1* was associated with Ezh2 and Hdac1 (Fig 3B). The RIP assay supported the potential of *malat1* in transcriptionally regulating gene expressions through its interaction with epigenetic modifiers. To explore this further, we checked the expression pattern of a few regeneration-associated genes (RAGs) on *malat1* knockdown. The RAGs such as *ascl1a*, *mmp9*, *lin28a,* and *wif1* were upregulated on *malat1* knockdown at 2dpi (Fig 3C). Analysis of a few other genes associated with cellular proliferation, such as *her4.1*, *insm1a,* and *egr1*, showed a *malat1*-MO dose-dependent decline in their expression (Fig 3D). A luciferase-based assay to validate *ascl1a, lin28a, mmp9, her4.1,* and *insm1a* promoter activity in zebrafish embryos revealed their dysregulation on *malat1* knockdown (Fig EV3A). These results supported the idea that *malat1* could be associated with epigenetic modifiers, thereby influencing gene expression. It is also interesting to note that several RAGs and epigenetic modifiers showed a decline in their protein levels in the *malat1* knockdown retina at 2 dpi (Fig EV3B), likely due to interference with mRNA translation by miRNAs like *let7*. Among these genes, *egr1*, a potent mitotic regulator, is located downstream of the *malat1* gene in zebrafish (Fig EV3C), indicating it may function in *cis* to mediate *malat1* influence on cellular proliferation during retina regeneration.

**Figure 3.**
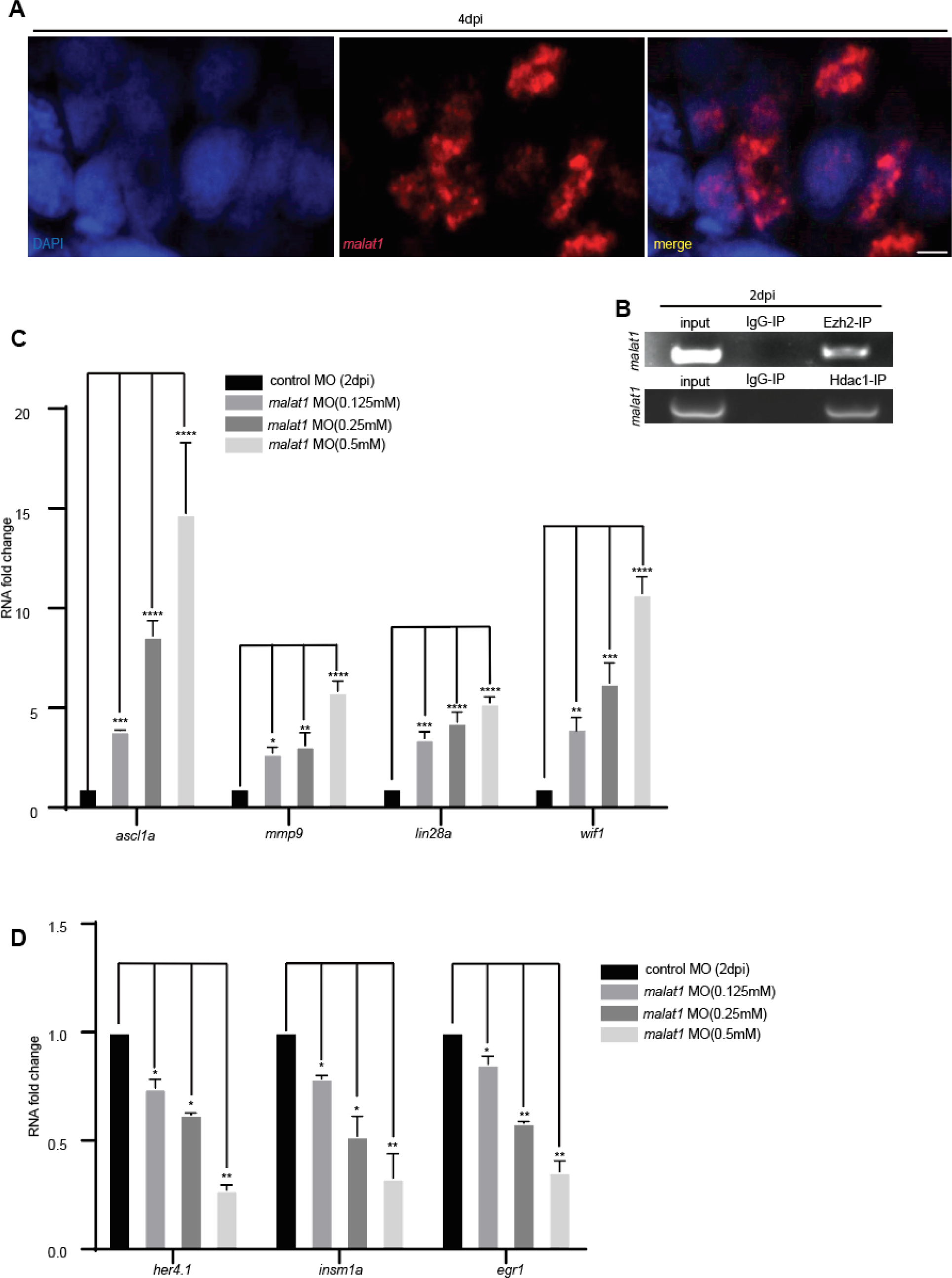
*malat1* localization and interaction with epigenetic modifiers to regulate regeneration-associated genes (RAGs). **(A)** Confocal images of fluorescence *in situ* hybridization (FISH) showing *malat1* localization in 4dpi retinal cross-sections after DAPI staining, indicating its presence at multiple nuclear loci. Scale bar: 10 µm. **(B)** *malat1* PCR check following RNA immunoprecipitation (RIP) in 2dpi retinal extract using anti-Ezh2 and anti-Hdac1 antibodies, with anti-IgG as a control, reveals a physical interaction between *malat1* and these epigenetic modifiers. **(C)** qPCR analysis of various regeneration-associated genes (RAGs) in *malat1* knockdown retinae at 2 days post-injury (dpi) shows increased expression of *ascl1a*, *mmp9*, *lin28a*, and *wif1.* Data is shown as mean ± standard deviation (SD), *p < 0.007; n = 6 biological replicates. **(D)** Conversely, a decrease in the expression of *her4.1*, *insm1a*, and *egr1* is observed in a concentration-dependent manner. Data is shown as mean ± standard deviation (SD), *p < 0.01; n = 6 biological replicates.

### Egr1 is a potential effector of *malat1* in inducing retinal progenitor proliferation

Given the syntenic conservation of many lncRNAs, we investigated genes near the *malat1* locus. Noting the proximity of *egr1* (Fig EV3C), we examined its role in retina regeneration and explored its temporal expression pattern post-retinal injury. Expression of *egr1* peaks at 16 hpi during Müller glia reprogramming and declines to below uninjured levels by 8 dpi (Fig EV4A). Knockdown of *egr1* in *1016tuba1a:GFP* transgenic retinas showed a dose-dependent reduction in progenitor cell numbers at 4 dpi (Fig EV4B and EV4C). Knockdown and overexpression of *egr1* resulted in decreased and increased *malat1* levels, respectively (Fig EV4D and EV4E), suggesting their interdependence during retina regeneration.

We hypothesized that the positive correlation between *malat1* and *egr1* expression plays a role in retinal progenitor proliferation. Transfecting injured retinas with *egr1* mRNA doubled progenitor proliferation. Notably, transfecting *egr1* mRNA alongside *malat1* MO rescued the decline in progenitor numbers caused by *malat1* knockdown at 4 dpi (Fig 4A and Fig 4B). These findings suggest that *malat1* influences retinal progenitors through Egr1, at least partially. qPCR analysis of cell-cycle regulators, including *cdk4*, *ccnd2*, and *cdkn1*, showed that Egr1 promotes cell cycle progression, accounting for the increased progenitor numbers in *egr1* mRNA-transfected retinas (Fig 4C).

**Figure 4.**
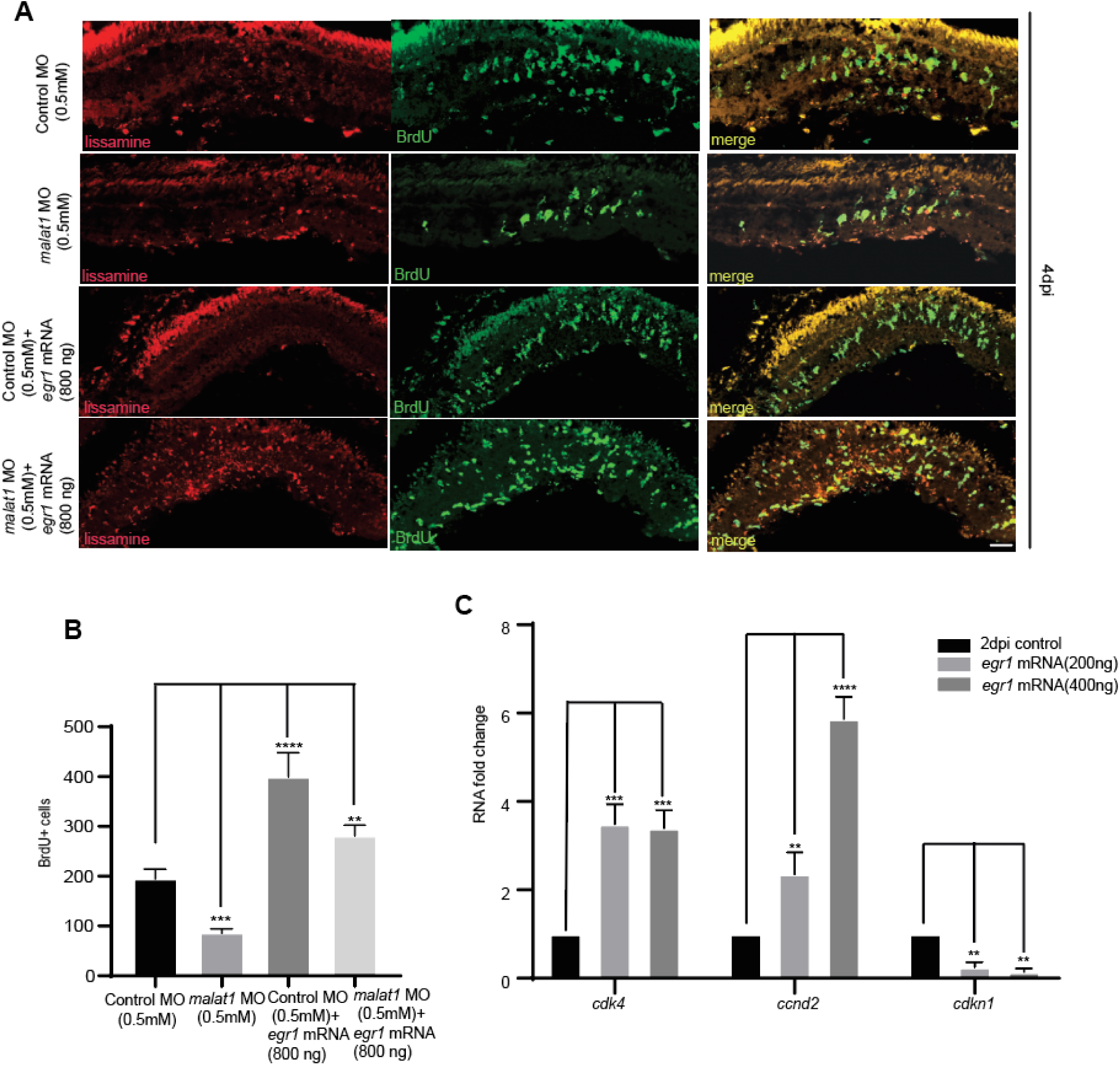
*malat1* knockdown effect is rescued by *egr1* overexpression and regulates cell cycle genes in MGPCs. **(A)** Confocal microscopy images show that the inhibitory effect of *malat1* knockdown on MGPC proliferation is reversed by overexpression of *egr1*, which is quantified in **(B)**. The data is presented as mean ± standard deviation (SD), *p < 0.0001; n = 6 biological replicates. This suggests that Egr1 partially mediates the activity of *malat1*. **(C)** qPCR analysis of cell cycle-related genes *cdk4*, *ccnd2*, and *cdkn1* in the context of *egr1* overexpression shows significant deregulation of these genes. These changes in gene expression led to alterations in the proliferative capacity of Müller glia-derived progenitor cells (MGPCs), highlighting the role of Egr1 in regulating cell cycle progression during retina regeneration. The data is presented as mean ± standard deviation (SD), *p < 0.05; n = 6 biological replicates.

### Egr1 positively influences retina regeneration through RAGs

Building on its effect on cell cycle progression, we explored if the *egr*1 could influence the regeneration-associated genes. The *egr1* mRNA transfected injured retina exhibited an enhanced expression of RAGs such as *ascl1a, mmp9, zic2b,* and *lin28a* (Fig 5A). The *egr1* mRNA transfection in the injured retina caused an abundance of retinal progenitor formation, traced up to 30 dpi, evidenced by the survival of proliferating cells marked with BrdU at 3^rd^, 4^th,^ and 5^th^ dpi. These progenitors could also differentiate into amacrine and bipolar retina cells (Fig 5B, 5C, and 5D). When transfected *in vivo* into zebrafish retina, *egr1* mRNA induced cell proliferation even in the uninjured retina (Fig EV5A and EV5B), but the induction of RAGs like *ascl1a, mmp9, zic2b, and lin28a* were minimal (Fig EV5C). BrdU-tracing of these proliferating cells showed no traceability by 30 days (Fig EV5D, EV5E, and EV5F), suggesting that while Egr1 promotes proliferation, injury-induced upregulation of RAGs is necessary for enough progenitor formation and its viability. This prompted us to investigate the cooperation between Egr1 and a crucial RAG, Ascl1a, to promote effective retina regeneration. To test this hypothesis, we explored the influence of *ascl1a* overexpression on progenitor proliferation in an injured retina, with or without *egr1* knockdown. Interestingly, even with *ascl1a* overexpression, we saw a decline in the retinal progenitor proliferation when *egr1* is downregulated (Fig 5E). Notably, despite a decrease in the retinal progenitor proliferation, the *ascl1a* overexpressed, and Egr1 declined retina had a higher number of retinal progenitors than that of *egr1* knockdown alone (Fig 5F). These findings suggest that Ascl1a and Egr1 influence retinal progenitor proliferation through parallel and synergistic routes.

**Figure 5.**
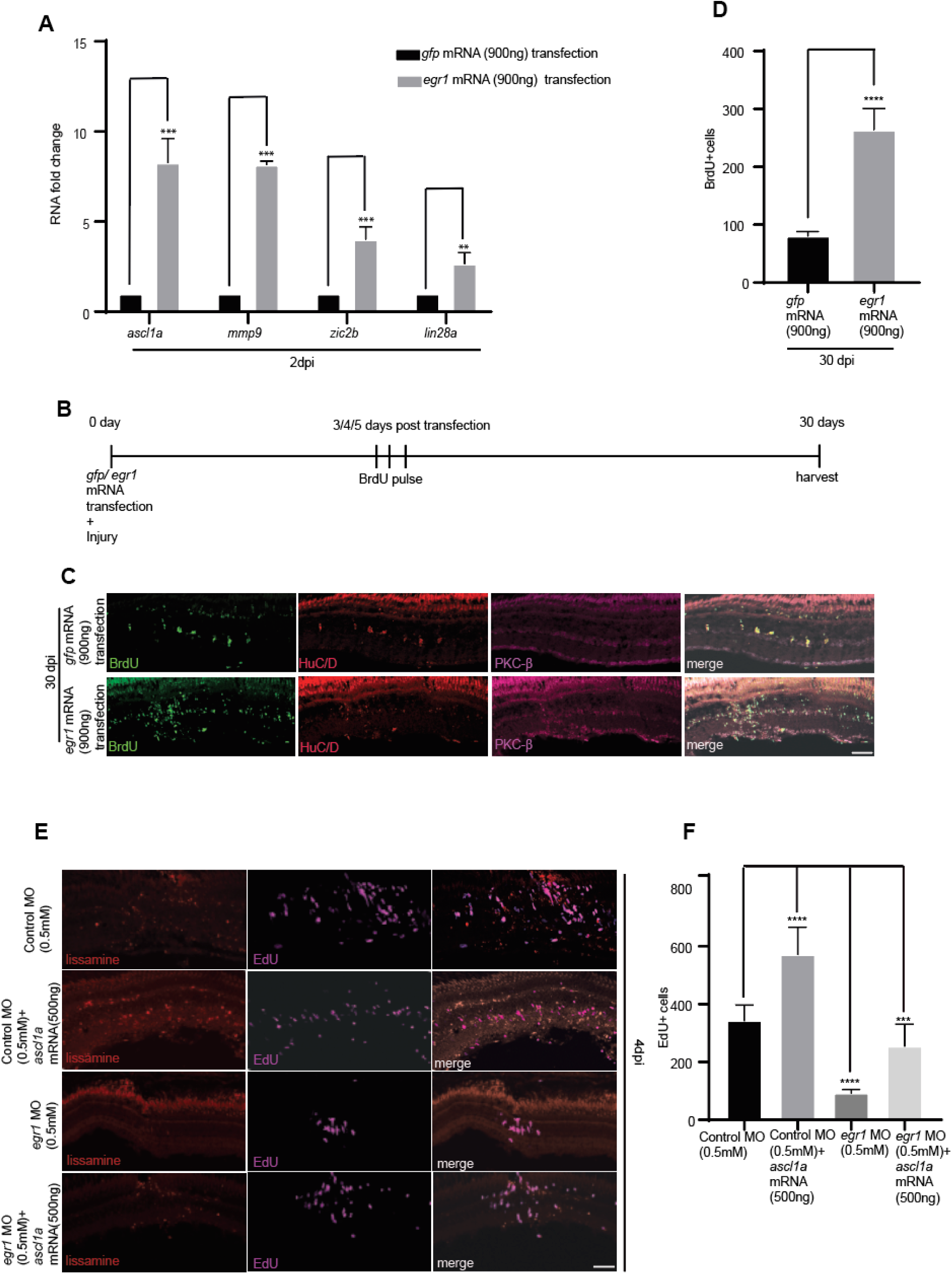
*egr1* overexpression enhances the proliferation and proliferated cells remained functional. Ascl1a requires Egr1 to fully mediate the regenerative response. **(A)** qPCR analysis at 2 dpi reveals that *egr1* overexpression enhanced the expression of regeneration-associated genes (RAGs) in response to retinal injury. Data is presented as mean ± standard deviation (SD), *p < 0.05; n = 6 biological replicates. **(B)** Schematic representation of the experimental design. **(C)** Confocal images of retinal cross-sections at 30 days post-injury (30 dpi) showing increased numbers of proliferated cells (BrdU +) following *egr1* overexpression compared to controls. Cells that incorporated BrdU at 3/4/5 dpi remained viable till 30 dpi. Quantification is provided in **(D)**. *p < 0.0005; n = 6 biological replicates. By 30 dpi, the proliferated cells (BrdU+) differentiated into HuD+ (amacrine cells) and PKC-β+ (bipolar cells), confirming they differentiated into different functional cell types. **(E)** Confocal images of retinal cross-sections demonstrating that knockdown of *egr1* on *ascl1a* overexpression decreases EdU+ cells compared to *ascl1a* overexpression alone, highlighting Egr1’s critical role in supporting the Ascl1a-driven regenerative response. Quantification is shown in **(F)**; *p < 0.0005; n = 6 biological replicates. Error bars represent standard deviation. Scale bars: 10 µm **(C, E)**.

### Egr1 is an effector of Wnt signaling in the regenerating retina

Wnt signaling is an important gene regulatory pathway that is shown to be essential during retina regeneration (Ramachandran *et al*, 2011). We were intrigued to explore whether Egr1 contributes as an effector of Wnt signaling in inducing retinal progenitors. We treated zebrafish retinas with SB216763, a GSK3β inhibitor that stabilizes β-catenin and activates Wnt signaling, with or without *egr1* MO, and observed its effect on proliferation at 4dpi (Fig 6A). SB216763 treatment increased retinal progenitors, but this effect was alleviated when *egr1* was knocked down (Fig 6A and 6B). The qPCR analysis of *egr1* mRNA in SB216763-treated retina revealed a dose-dependent increase in its expression levels at 2dpi (Fig 6C). However, the SB216763 drug treatment caused a decrease in the *malat1* expression (Fig 6D), which suggested the possible direct regulation of *egr1* mRNA through Wnt signaling. To test this possibility, we analyzed the regulatory DNA elements of the *egr1* promoter for potential TCF/LEF binding sites. We identified three TCF/LEF binding sites immediately upstream of the *egr1* gene (Fig 6E). A ChIP assay at 2dpi using a β-catenin antibody showed occupancy of β-catenin to these sites on the *egr1* promoter (Fig 6F), supporting that Wnt signaling directly regulates Egr1 expression during retina regeneration.

**Figure 6.**
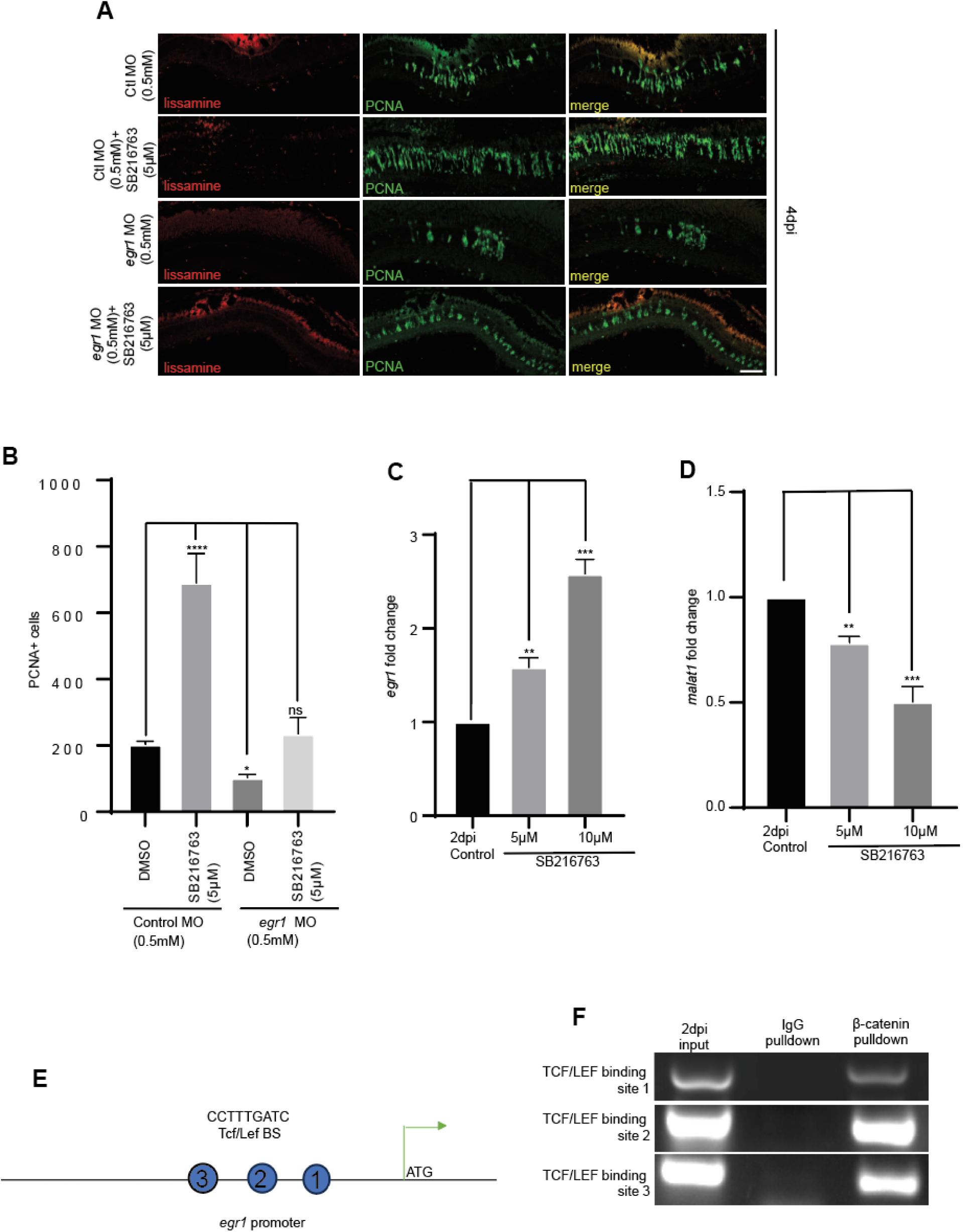
Wnt signaling mediates its effect by inducing *egr1*. **(A)** Confocal images of retinal cross-sections reveal that the increase in PCNA+ cells induced by β-catenin stabilization through SB216763 drug administration was reduced following egr1 knockdown. It has been quantified in **(B).** It hints that Wnt signaling may partly regulate MGPC proliferation by modulating *egr1* levels. The data is presented as mean ± standard deviation (SD), *p < 0.0003; n = 6 biological replicates. Scale bars: 10 µm. **(C)** qPCR analysis demonstrates that *egr1* expression is significantly induced by stabilizing β-catenin (SB216763 administration), indicating that Wnt signaling plays a crucial role in regulating *egr1* expression during retinal regeneration. The data is presented as mean ± standard deviation (SD), *p < 0.05; n = 6 biological replicates. **(D)** qPCR showing *malat1* levels get reduced on stabilizing β-catenin (SB216763 administration), *p < 0.01; n = 6 biological replicates. **(E)** Schematic representation of the TCF/LEF binding site (β-catenin binding site) on the *egr1* promoter. **(F)** PCR analysis following ChIP with anti-β-catenin antibody confirms that β-catenin directly binds to the *egr1* promoter to drive its transcription.

To further verify it, we treated injured retinas with XAV939, a Wnt signaling inhibitor, with or without *egr1* overexpression (Fig EV6A). While XAV939 treatment reduced retinal progenitor numbers, *egr1* overexpression rescued the progenitor count to control levels (Fig EV6A and EV6B). These observations suggest that Wnt signaling influences the induction of retinal progenitors through Egr1 and that the blockade of Wnt signaling could be rescued through *egr1* overexpression. The qPCR analysis of the *egr1* mRNA in SB216763 and XAV939 treated conditions reveals a positive interplay between Wnt signaling and Egr1 (Fig 6C and EV6C). This suggests that Wnt signaling regulates retinal progenitor induction through Egr1.

### *malat1* is suppressed by Delta-Notch signaling and functions to induce deltaD

Previous studies have characterized the importance of Delta-Notch signaling in zebrafish retina regeneration, which impinges on expanding the proliferation zone. Negative regulation of Delta-Notch signaling expanded the regenerative zone, as indicated by increased proliferating retinal progenitors. The Delta-Notch signaling is mediated through the Notch intracellular domain (NICD), which induces a transcriptional repressor hairy enhancer of split (Her4.1) (Campbell *et al*, 2022). We investigated whether Her4.1 affects the *malat1* expression. For this, we either downregulated or overexpressed *her4.1* in the injured retina and checked for the *malat1* levels (Fig 7A). Knockdown of *her4.1* increased (Fig 7B and 7C), while its overexpression reduced the *malat1* expression (Fig 7B and 7D). Notably, the seclusion of *malat1* from the proliferating cells observed in the normally regenerating retina was also disrupted, with *malat1* also being expressed in proliferating cells in the *her4.1* downregulated (Fig 7E and 7F). We further explored the *malat1* promoter activity in the presence of *her4.1*-targeting MO or its mRNA using luciferase assay in zebrafish embryos. We saw a negative regulation of the *malat1* promoter by Her4.1 (Fig EV7A). These results suggest that the regulation of *malat1* is mediated through Delta-Notch signaling mainly to restrict its expression to the neighboring cells of the proliferating progenitors.

**Figure 7.**
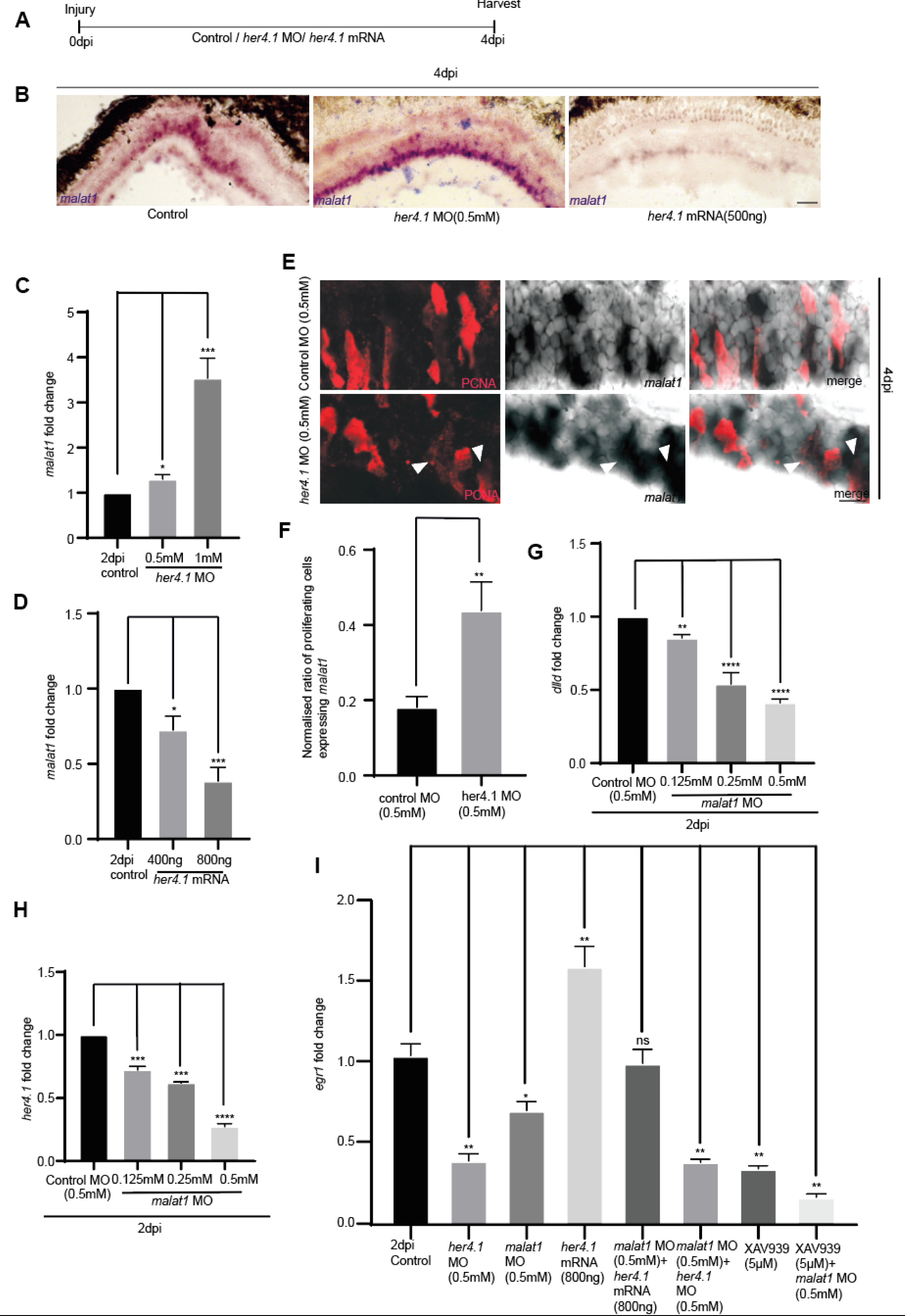
*malat1* regulates *egr1* in a cell non-autonomous manner via Notch signaling. Wnt signaling and Notch signaling synergistically regulate *egr1* levels. **(A)** Experimental timeline schematic **(B)** Bright-field image of retinal cross-sections showing *malat1 in situ* hybridization after *her4.1* knockdown and overexpression respectively, compared to control. **(C)** qPCR analysis of *malat1* levels on *her 4.1* knockdown, *p < 0.01; n = 6 biological replicates, and **(D)** *her 4.1* overexpression, *p < 0.03; n = 6 biological replicates. The data is represented as mean ± standard deviation (SD). **(E)** 60X confocal images showing *malat1* expression gets induced in proliferating cells after *her4.1* knockdown, quantified in **(F).** The data is presented as mean ± standard deviation (SD), *p < 0.003; n = 6 biological replicates. Arrowheads indicate *malat1* localization in proliferating cells. This observation suggests that *her4.1* suppresses *malat1* expression in these cells. **(G)** qPCR analysis showing the impact of *malat1* knockdown on *dlld* expression, *p < 0.003; n = 6 biological replicates, and *her4.1* expression **(H)**, *p < 0.005; n = 6 biological replicates. The data is presented as mean ± standard deviation (SD). **(I)** qPCR analysis shows that combined knockdown of *her4.1* and *malat1* reduces *egr1* levels to the same extent as *her4.1* knockdown alone, suggesting that *malat1* regulates *egr1* through *her4.1* (Notch signaling). Combined inhibition of Wnt signaling (XAV939) and *malat1* knockdown caused an even greater decrease in *egr1* expression. This suggests that *malat1* regulates *egr1* levels in proliferating cells through Notch signaling, while Notch and Wnt signaling pathways independently regulate *egr1* levels. The data is represented as mean ± standard deviation (SD), *p < 0.005; n = 6 biological replicates. Scale bars represent 10 µm in **(B)** and **(E)**.

Fluorescence *in situ* hybridization (FISH) was conducted using antisense probes for *malat1*, *dlld* (DeltaD encoding transcript), and *her4.1* mRNA. Co-labeling of *malat1* and *dlld* was observed (Fig EV7B), while *malat1* and *her4.1* showed mutual exclusion in the retina at 4 dpi (Fig EV7C). A cell-sorting analysis of *1016tuba1a*:GFP transgenic retina at 4dpi confirmed the abundance of *dlld* in GFP- and *her4.1* expression in GFP^+^ cells (Fig EV7D). The *her4.1* expression is a feature of proliferating retinal progenitors (Mitra *et al*, 2018). These findings prompted us to explore whether *malat1* expression in the neighboring cells influenced Delta-Notch signaling. We speculated if the *malat1* could influence the expression of Delta ligand, leading to the activation of Notch receptors in the proliferating retinal progenitors. The *malat1* knockdown caused a decline in the *dlld* mRNA in a dose-dependent manner at 2dpi (Fig 7G). This decline in the *dlld* was also accompanied by a decrease in the *her4.1* mRNA levels (Fig 7H). This suggests that *malat1* regulates Notch signaling by affecting *dlld* expression, eventually influencing *her4.1* levels in proliferating cells.

Initially, we hypothesized that *malat1* regulated *egr1* in a *cis* manner, implying that both should be expressed in the same cell. However, given that Wnt signaling is active in proliferating cells (Meyers et al., 2012; Ramachandran et al., 2011) and promotes *egr1* expression, which pushes cells to enter into the cycling phase, we reassessed the localization of *egr1*. Cell-sorting analysis of *1016tuba1a*:GFP transgenic retina at 4dpi confirmed the abundance of *egr1* in GFP+ cells (Fig EV7D). It suggests that *malat1* may regulate *egr1* non cell-autonomously, potentially through juxtacrine signaling mechanisms like Notch signaling. Knockdown experiments demonstrated that both *her4.1* and *malat1* downregulate *egr1*, but the effect of combined knockdown mirrored that of *her4.1* knockdown alone, suggesting that *malat1* acts on *egr1* primarily through *her4.1*. Additionally, inhibiting Wnt signaling using XAV939 alongside *malat1* knockdown produced a synergistic reduction in *egr1* (Fig 7I), highlighting the independent contributions of these pathways in inducing retinal progenitors.

### TGF-β signaling suppresses stable *malat1* expression in zebrafish but enhances *Malat1* expression in mice

Tgf-β signaling is one of the major pro-proliferative pathways essential for zebrafish retina regeneration (Sharma *et al*, 2020). We explored whether Tgf-β signaling plays a role in regulating *malat1* expression. Inhibition of Tgf-β signaling in the regenerating retina using the blocker SB431542 caused an elevated expression of *malat1* in a dose-dependent manner (Fig 8A). Conversely, the Tgf-β1 protein inhibited *malat1* expression (Fig 8A). The *malat1* RNA *in situ* hybridization performed in 4dpi retina (Fig 8B) and the luciferase assay performed in zebrafish embryos using *malat1* promoter constructs (Fig EV8A) confirmed these results. A closer look at the regulatory DNA elements of the *malat1* gene revealed two 5GC elements, which are putative pSmad3 binding sites essential for transcriptional activation by Tgf-β signaling (Martin-Malpartida *et al*, 2017)et al. (Fig 8C). A ChIP assay performed in 2dpi retinal protein extracts using the pSmad3 antibody confirmed that pSmad3 occupied these 5GC elements (Fig 8D). This seemed a conundrum, as qPCR, RNA *in situ* hybridization, and luciferase assay results proved that Tgf-β signaling is a negative regulator of *malat1* gene expression.

**Figure 8.**
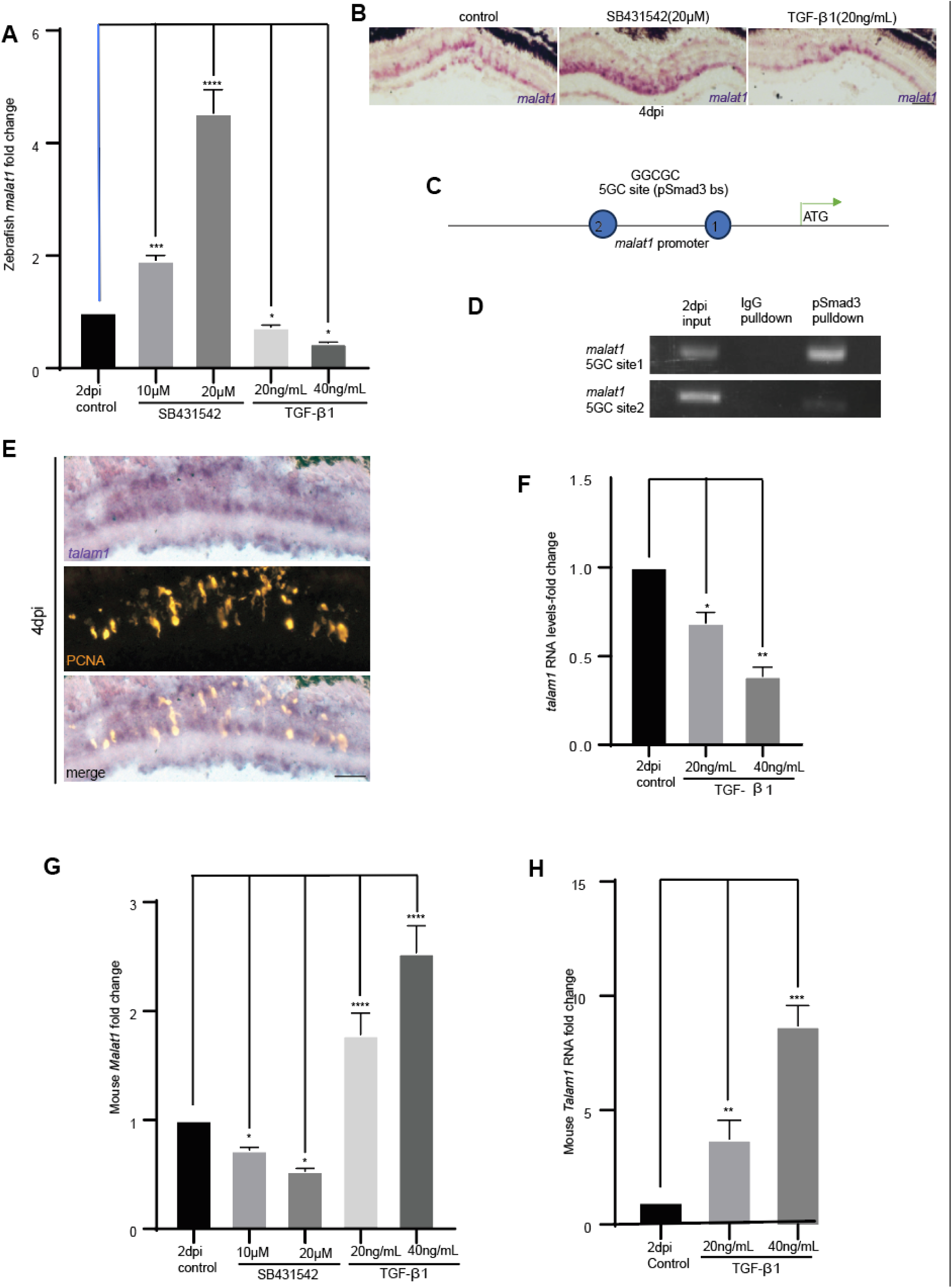
Differential regulation of *malat1* by TGF-β signaling in zebrafish and mice, and the interplay of *talam1* in mediating this regulation. **(A)** qPCR analysis of *malat1* levels following inhibition and overexpression of TGF-β signaling in zebrafish retinae. The data is presented as mean ± standard deviation (SD), *p < 0.05; n = 6 biological replicates. **(B)** Bright-field images of retinal cross-sections following *malat1 in situ* hybridization in zebrafish treated with SB431542 (TGF-β inhibitor) and in TGF-β1 overexpression scenarios, revealing its repressive role on *malat1*expression. **(C)** Diagrammatic representation of 5GC elements on the *malat1* promoter in zebrafish, illustrating the potential regulatory sites for TGF-β signaling**. (D)** PCR analysis after pSmad3 ChIP in 2dpi zebrafish retina shows that pSmad3 occupies 5GC elements on the zebrafish *malat1* promoter. IgG pulldown serves as a negative control, revealing the occupancy of pSmad3 on 5GC elements of the *malat1* promoter. **(E)** 20X bright-field (BF) images of retinal cross-sections showing distinct *talam1* expression after *in situ* hybridization using a probe specific for *talam1*. **(F)** ss-qPCR analysis in TGF-β1 overexpression scenarios showing downregulation of *talam1*, indicating the inhibitory role of TGF-β signaling on *talam1* in zebrafish retina regeneration. The data is presented as mean ± standard deviation (SD), *p < 0.05; n = 6 biological replicates. **(G)** qPCR analysis of mouse *Malat1* levels following inhibition and overexpression of TGF-β signaling in injured mouse retinae. The data is presented as mean ± standard deviation (SD), *p < 0.04; n = 6 biological replicates. **(H)** ss-qPCR analysis showing that *Talam1* is upregulated in the injured mouse retina following TGF-β1 overexpression. The data is presented as mean ± standard deviation (SD), *p < 0.05; n = 6 biological replicates. The scale bar is 10 µm in **(B)** and **(E)**.

Previous studies have indicated that *talam1*, an antisense transcript of *malat1*, is essential for the maturation of the *malat1* transcript. Mature *malat1* has improved half-life and is functional. (Gomes *et al*, 2019; Zong *et al*, 2016). The genomic location of both genes is on sense and antisense strands of the gene loci (Fig EV8B). The RNA-*in situ* hybridization also confirmed the expression of *talam1* in 4dpi retina (Fig 8E). The promoter of *talam1* harbored pSmad3-binding TIE elements (Fig EV8C). TIE elements are sites wherein the pSmad3 binds with other protein factors to cause transcriptional repression (Kerr *et al*, 1990; Sharma *et al*., 2020). A ChIP assay performed in 2dpi retinal protein extracts using pSmad3 antibody confirmed the occupancy of these TIE elements of *talam1* promoter by pSmad3 (Fig EV8D). Tgf-β1 protein delivery caused a decline in the expression levels of *talam1* in the 2dpi retina (Fig 8F). This decline of *talam1* levels could probably cause reduced mature and functional *malat1* transcript levels in Tgf-β signaling-induced zebrafish retina.

In mammals, the Tgf-β signaling is anti-proliferative except in cancerous conditions (Close *et al*, 2005). We were intrigued to explore whether the *malat1* is differentially regulated during mice retina regeneration compared to zebrafish. The mouse *Malat1* gene promoter analysis revealed the presence of both 5GC and TIE elements (Fig EV8E). A ChIP assay performed using PSmad3 antibody in retinal protein extracts of 2dpi mice retina revealed that, of the five 5GC sites, only two of them were occupied by PSmad3, while all the three TIE elements were bound by PSmad3 (Fig EV8F). Contrary to the zebrafish retina, inhibiting TGF-β signaling in the injured mice retina at 2dpi downregulated *Malat1* RNA levels (Fig 8G). The TGF-β1 protein caused an upregulation of both *Malat1* (Fig 8G) and *Talam1* (Fig 8H) transcript levels in mice retinas at 2dpi. We also found an anticipated decrease of *egr1* mRNA levels in zebrafish (Fig EV8G) and an increase in *Egr1* mRNA levels in mice retina (Fig EV8H) because of TGF-β1 protein delivery.

We performed a strand-specific RT-PCR of *talam1* in zebrafish embryos to validate its expression during embryonic development (Fig EV8I). Having found its expression in zebrafish embryos, we created transgenic lines of zebrafish driving GFP under the control of *talam1* (Fig EV8J) and *malat1* promoters (Fig EV8K). Both these lines showed overlapping yet differential expression of GFP, suggesting that the interplay of *talam1* and *malat1* occurs during embryonic development. Together, these findings support the view that the differential regulation of *malat1* in mice and zebrafish through TGF-β signaling could also account for the lack of retinal regeneration in mice compared to zebrafish.

Inhibition of TGF-β signaling in zebrafish reduced proliferation but paradoxically increased *egr1*, a pro-proliferative factor, expression (Fig EV8G). After the retinal injury, manipulating TGF-β signaling and Egr1 expression revealed that co-overexpression of both caused a synergistic decrease in proliferation (Fig 9A and 9B). Co-overexpression of TGF-β1 and *egr1* significantly elevated *tbx2a* levels, a pro-proliferative factor (Fig 9C). Based on previous studies (Mohamad *et al*, 2018), there is possible co-sequestration of Tbx2a and Egr1, negating both factors’ pro-proliferative effects. Decreased *cdk4* and increased *cdkn1* levels in the scenario supported the hypothesis that Egr1 cannot perform its function in the co-overexpression scenario (Fig 9D). In mice, TGF-β signaling upregulates Egr1, similar to the co-overexpression scenario observed in zebrafish, potentially explaining the opposing effects of TGF-β signaling on proliferation. A schematic of the gene regulatory network elucidated in this study is depicted (Figure EV9).

**Figure 9.**
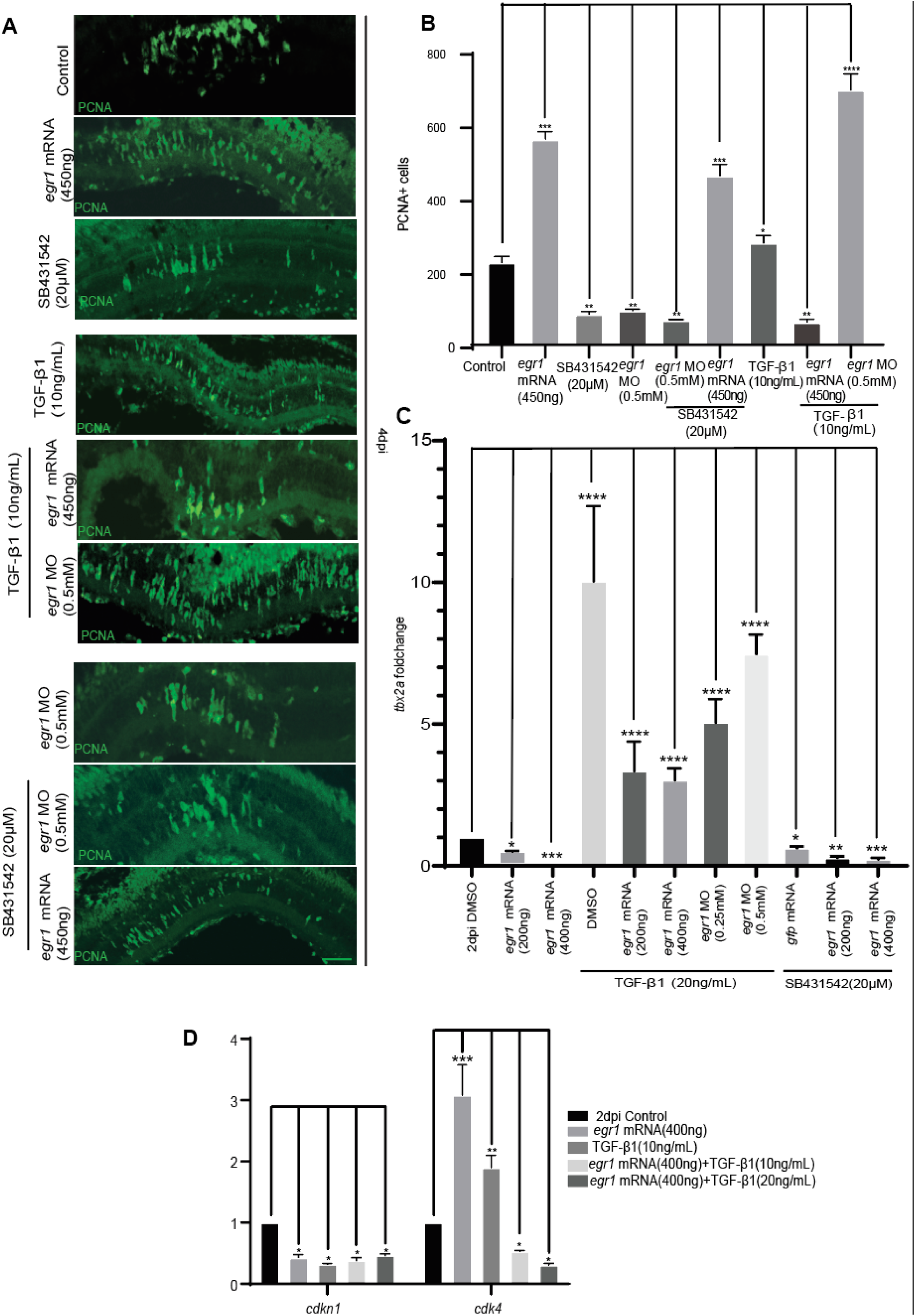
Impact of TGF-β signaling and Egr1 on proliferation and cell cycle regulation during retina regeneration. **(A)** Confocal images of retinal cross-sections at 4 days post-injury (4 dpi) showing proliferation status upon modulating TGF-β signaling and Egr1 levels, individually and in combination. Scale bar: 10 µm. Quantification is shown in **(B)**. The data reveal that TGF-β signaling and Egr1 individually promote proliferation, but their combined overexpression decreases PCNA+ cells. Data is presented as mean ± standard deviation (SD), *p < 0.005; n = 6 biological replicates. **(C)** qPCR analysis of *tbx2a* expression at 2 dpi under conditions of TGF-β1 overexpression with or without *egr1* overexpression or knockdown, and TGF-β signaling inhibition (SB431542), alone or combined with *egr1* overexpression or knockdown. The data is presented as mean ± standard deviation (SD), *p < 0.05; n = 6 biological replicates. **(D)** qPCR analysis shows synergistically reduced *cdk4* expression levels and increased *cdkn1* expression when TGF-β1 and Egr1 are overexpressed in combination. This suggests that Egr1 may be sequestered away from its target sites under these conditions, altering cell cycle regulation. Data is presented as mean ± standard deviation (SD), *p < 0.05; n = 6 biological replicates. The scale bar is 10 µm in **(A)**.

## Discussion

Retina regeneration is bliss in lower vertebrates such as fishes and frogs. However, mammals do not have this ability, especially in central nervous system organs such as the retina. Manipulating gene expression pathways in mouse retina that emulate zebrafish retina regeneration has mostly resulted in positive responses, albeit incomplete. Several holistic gene expression studies have revealed that the mammalian retina exhibits some regenerative potential, though it remains incomplete. In this study, we tried to explore the importance of long non-coding RNA *malat1* during retina regeneration. We also explored if *malat1* is differentially regulated in mice and zebrafish retina, which could probably account for the lack of complete regeneration in injured mice retina. *Malat1* has been extensively studied to contribute to several biological phenomena, including wound healing (Chen *et al*, 2023; Liu *et al*, 2019), cancer initiation, and metastasis (Arun *et al*, 2020; Hussein *et al*, 2024; Liu *et al*, 2017). *Malat1* is known to promote cellular proliferation and is considered to be a curative target for cancer (Amodio *et al*, 2018; Bhat *et al*, 2024) and metabolic syndromes such as type-2 diabetes (Aihemaiti *et al*, 2024). The roles of *malat1* are explored and demonstrated in bone fracture healing (Qin *et al*, 2024), peripheral nervous system repair (Wu *et al*, 2020), and muscle (Chen *et al*., 2017) and endothelium (Chen *et al*, 2021) regeneration. Hence, we anticipated the contribution of *malat1* to retina regeneration.

At first, we explored the importance of *malat1* during zebrafish development, as no phenotype was observed in mice after *Malat1* knock-out (Zhang *et al*, 2012). We saw a significant mortality of zebrafish embryos because of *malat1* knockdown, suggesting important gene expression events are guided through this lncRNA, unlike in mice. It is to be noted that despite normal development, the *Malat1* knock-out disrupted many adjacent genes of the *Malat1* locus (Zhang *et al*., 2012). A similar phenomenon could also be at play in zebrafish, as we found a significant decline in the retinal progenitor formation upon *malat1* knockdown in zebrafish retina at early and late time points.

The *malat1* RNA is nuclear localized in its expression, which indicates its genomic functions. The *Malat1* is shown to interact with nuclear proteins such as Nucleolin and Nucleophosmin (Ghosh *et al*, 2023). The forced expulsion of *Malat1* from the nucleus causes dysregulation of TDP-43, whose aggregation is implied in neurodegeneration (Nguyen *et al*, 2020). Our study also reveals the interaction of *malat1* to histone deacetylase Hdac1 and histone methylase Ezh2 of polycomb repressor complex 2 in zebrafish retina during regeneration. These findings suggest the involvement of *malat1* in the epigenetic regulation of various chromatin loci. Although nuclear, a portion of the cytoplasmic localization of *Malat1* in the brain serves as a translatable coding RNA by synaptic stimulation (Xiao *et al*, 2024).

*Malat1* is known for its role in diverse cell types and influence on other signaling pathways such as Hippo-Yap and Wnt signaling. The *MALAT1* level stays upregulated in diabetic cardiomyopathy. The knockdown of *MALAT1* in the diabetic cardiomyopathy mice model alleviates the clinical manifestations of high glucose on heart muscles, such as inflammation and collagen accumulation, by influencing the Hippo-Yap pathway (Liu *et al*, 2020). *Malat1* is regulated epigenetically through histone demethylase JMJD2C at H3K9me3 and H3K36me3 locations of the *Malat1* promoter, causing its upregulation and subsequent activation of Wnt signaling in cancer metastasis (Wu *et al*, 2019b). These scenarios prompted us to explore the expression dynamics of *malat1* during retina regeneration and its implications in downstream gene expressions and signaling pathways.

The physical proximity of one of the pro-mitotic genes, *egr1,* to the *malat1* locus in linear DNA was unique compared to human and mouse loci. When exploring *egr1* levels in *malat1* knockdown, we saw a dose-dependent decrease in *egr1* mRNA. The knockdown of *egr1* also caused a decline in retinal progenitors. Interestingly, the declining retinal progenitor number because of the *malat1* knockdown could be rescued by *egr1* overexpression. This finding suggests that one of the regulatory pathways governed by *malat1* is via Egr1. It is also important to note that several RAGs, such as *ascl1a*, *sox2*, *lin28a*, *hdac1*, and *yap1,* were downregulated because of *malat1* knockdown. It was intriguing to note that the pSmad3 protein, an effector of Tgf-β signaling, was upregulated in the *malat1* knockdown retina. This observation was indicative of an interrelationship between Tgf-β signaling and *malat1*.

The Retinal progenitor proliferation is strongly influenced by Wnt signaling during retina regeneration in zebrafish (Ramachandran *et al*., 2011). The Wnt signaling could also influence mammalian retinal neuronal reprogramming and regeneration (Osakada *et al*, 2007; Sanges *et al*, 2013). Pharmacological stabilization of β-catenin enhanced retinal progenitor formation in the injured retina and could facilitate retinal progenitor formation in the uninjured retina in zebrafish (Ramachandran *et al*., 2011). The SB216763 drug, when used to stabilize β-catenin, caused an upregulation of *egr1* and enhanced retinal progenitor number. This increase of retinal progenitors was nullified by *egr1* knockdown. The *egr1* promoter had several TCF/LEF binding sites favoring β-catenin binding for transactivation. Negative regulation of Wnt signaling using the XAV939 drug caused a decline in retinal progenitor but enhanced *malat1* levels. However, this reduction in retinal progenitors because of XAV939 could be nullified by *egr1* overexpression. Thus, Egr1 could be an important effector of Wnt signaling in stimulating proliferation during retina regeneration in zebrafish.

*egr1* mRNA is expressed in retinal progenitors, while *malat1* lncRNA is restricted to neighboring cells. The reduction of *egr1* levels upon *malat1* knockdown in the retina suggests Delta-Notch signaling mediates their interplay, prompting us to explore its role in this regulation. Blocking the Delta-Notch signaling effector Her4.1 results in a reduction of retinal progenitors. Being a transcriptional repressor, we speculated if Her4.1 negatively regulated *malat1* expression. We confirmed that Her4.1 negatively affected *malat1* transcription. More importantly, *malat1* did not remain excluded from the sparsely present proliferating cells in the *her4.1* knockdown retina.

Interfering with the Tgf-β signaling had a profound effect on *malat1* levels. *Malat1* promoter activity decreased with active Tgf-β signaling but increased with SB431542 treatment. Despite containing the pSmad3 transactivation domain, the 5GC element, the *malat1* promoter appeared ineffective. *malat1* is known to be regulated via the antisense transcript *talam1,* whose presence dictates the stability and functionality of *malat1* RNA (Gomes *et al*., 2019). Interestingly, the zebrafish *talam1* promoter had several TIE elements bound by pSmad3, thus contributing to its transcriptional repression. This inhibition of *talam1* by Tgf-β signaling could be crucial in the *malat1* dynamics. A reduction in egr1 mRNA expression also mirrors the decrease in malat1 levels. However, it is to be noted that despite a decline in *talam1*, mature *malat1,* and *egr1* in active Tgf-β signaling, it did not cause a decline in the retinal progenitor as expected with *malat1* knockdown. This could be possible because of many other gene expression events during the early and late phases of retina regeneration, regulated during retina regeneration through Tgf-β signaling (Sharma *et al*., 2020). Presumably, the downregulation of *talam1*, *malat1,* and *egr1* in activated Tgf-β signaling played a balancing role in restricting the retinal progenitor number within optimal range, as done by Delta-Notch signaling in regenerating zebrafish retina (Wan *et al*, 2012).

Exploration of TGF-β signaling during mice retina regeneration demonstrated that *Malat1* and *Talam1* levels are upregulated because of TGF-β1 protein delivery. The *Egr1* levels also get positively regulated in a TGF-β1 protein concentration-dependent manner in mice retina. However, being anti-proliferative, the TGF-β signaling in mammals did not cause retinal progenitor proliferation, unlike zebrafish (Sharma *et al*., 2020; Zhang *et al*, 2017). In this study, we demonstrated that TGF-β signaling inhibited *egr1* expression in zebrafish, a finding that contrasts with observations in mice, where TGF-β signaling positively regulates *Egr1*. In zebrafish, blocking TGF-β signaling led to upregulating the pro-proliferative gene *egr1*, which did not increase retinal progenitors. Surprisingly, co-overexpression of TGF-β1 and *egr1* caused a decline in progenitor cell numbers. Various reports show that TBX2 protein interacts with Egr1 and sequesters it away from its target sites (Crawford *et al*, 2019; Mohamad *et al*., 2018). We saw that TGF-β signaling induced *tbx2a* expression, a pro-proliferative factor, while *egr1* downregulated it. The elevated levels of TGF-β1 and *egr1* significantly upregulated *tbx2a* levels, suggesting the co-sequestration of Tbx2a and Egr1, nullifying the pro-proliferative effect of each other. In mice, TGF-β signaling positively regulated *Egr1,* possibly via *Malat1*, but their simultaneous elevation could inhibit *Egr1*’s pro-proliferative function due to *Tbx2* interplay. This finding highlights how the same pathway can produce opposing effects on proliferation depending on species and context. Modulating Egr1 during active TGF-β signaling may offer a therapeutic strategy to enhance mammalian retinal regeneration.

## Materials and methods

### Zebrafish husbandry, retinal injury, and pharmacological agents

In circulating aquatic systems, wild-type zebrafish (*Danio rerio*) were maintained at 26–28.5°C on a 14:10-hour light/dark cycle. Embryos were obtained by natural breeding. Transgenic *1016 tuba1a: GFP* zebrafish (Fausett & Goldman, 2006) were also used in this study. Retinal injuries were induced in 8–12-month-old fish under tricaine methane sulfonate anesthesia with a 30G needle (Fausett & Goldman, 2006; Sharma & Ramachandran, 2019).

C57BL/6 mice were housed on a 12-hour light/dark cycle with *ad libitum* food and water. The retinal injury was induced, under isoflurane anesthesia, by injecting 100 mM NMDA intravitreally. All procedures followed institutional ethical guidelines and were IAEC-approved.

Unless otherwise specified, drug and protein injections into the vitreous were performed during the injury process, using a 5 µL Hamilton syringe, equipped with a 30G needle. All pharmacological agents used namely SB431542, TGF-β inhibitor (Sigma Aldrich, S4317), XAV939, Wnt signaling inhibitor (Tocris), glycogen synthase kinase-3β (GSK-3β) inhibitor I (SB216763, EMD Calbiochem), were dissolved in DMSO, to make stock concentration as 1mM, which was diluted further in PBS or autoclaved de-ionized water. TGF-β1 protein (Abcam, ab50036) was prepared at 4 mg/mL in 10 mM citric acid, with working concentrations diluted in PBS or autoclaved deionized water. Chemical-mediated injury with NMDA, N-Methyl D-aspartic acid (sigma, M3262), was performed by delivering it to vitreous humor in zebrafish and mice eyes, through the cornea.

### Transgenic line generation

Transgenic lines were generated using the Tol2 system with *pT2AL200R150G* vector (Urasaki *et al*, 2006). ∼4 kb *malat1* and *talam1* promoters were cloned upstream of GFP in the vector and the purified construct was injected with 25 ng *tol2* mRNA into one-cell zebrafish embryos, then screened for GFP.

### Antisense morpholino transfection, *in vivo* mRNA transfection

For gene knockdown experiments, approximately 0.5μl of Lissamine-tagged morpholinos (MOs) (Gene Tools) at a concentration of 0.125 to 1.0 mM were administered during retinal injury using a 5 μl Hamilton syringe. To facilitate MO delivery into retinal cells, electroporation was performed following the method described by (Fausett *et al*, 2008). Sequences of morpholinos used in the study are:

Control MO: 5′CCTCTTACCTCAGTTACAATTTATA-3′ (Fausett *et al*., 2008)

*ascl1a* MO: 5′ATCTTGGCGGTGATGTCCATTTCGC-3′ (Fausett *et al*., 2008)

*malat1* MO: 5′CCACCAGGGTCTTTTGCTTTTTTTC-3′ (Wu *et al*., 2019a)

*egr1* MO: 5’-GCA GCC ATC TCT CTG GAG TGT GCT C-3′ (Hu *et al*, 2006)

*her4.1* MO: 5’-TTGATCCAGTGATTGTAGGAGTCAT-3’ (Mitra *et al*., 2018)

For *in-vivo* mRNA transfection, the pCS2 vector harboring the full-length coding sequence of genes, in correct orientation was linearised using NotI/KpnI, and capped mRNA was synthesized *in-vitro* using mMESSAGE mMACHINE kit (Ambion), as per manufacturer’s instructions. A mix containing lipofectamine, HBSS, and mRNA was prepared as described in (Mitra *et al*., 2018; Sharma *et al*, 2019; Sharma & Ramachandran, 2019). The mix was delivered intravitreally through the cornea using a Hamilton syringe. The transfection mixture containing *gfp* mRNA was parallelly injected into the control retina. For embryos, ∼200nL of morpholino or mRNA mix was injected in 1 cell stage.

### Primers and plasmid construction

Table I lists all the primers used in the study for qPCR, cloning, SDM (site-directed mutagenesis), and probe template. The coding sequence of *malat1* and *egr1* were amplified from 24 hpf zebrafish embryos, and cloned in pCS2+ vector, using respective restriction sites. The *ascl1a:*GFP*-luciferase, lin28a:*GFP*-luciferase, insm1a:*GFP*-luciferase*, *her4.1:*GFP*-luciferase* and *mmp9:*GFP*-luciferase* constructs have been described previously (Ramachandran *et al*, 2010) (Kaur *et al*, 2018; Ramachandran *et al*, 2012). *malat1* and *talam1* promoter (∼4kb) was amplified from zebrafish genomic DNA, and cloned into *pT2AL200R150G* vector, upstream of *gfp*, using primers harboring XhoI and SalI sites. *malat1* promoter was also cloned into the pEL luciferase expression vector using primers harboring XhoI and EcoRI sites, to generate *malat1:*GFP*-luciferase* construct.

Site-directed mutagenesis was performed to add *malat1* morpholino binding site, upstream of GFP, on the pEGFP-N1 construct, using the protocol described in (Ramachandran *et al*., 2012). Templates for the *dlld* probe were amplified from 24 hpf embryos using primer pairs with a T3 polymerase binding site on the reverse primer, as listed in Table I. The *talam1* probe was designed by cloning the *malat1* splice site region into the pCS2+ vector using ClaI/StuI restriction sites, linearized with ClaI, and transcribed using SP6 polymerase.

### ChIP assay

Chromatin immunoprecipitation (ChIP) assays were conducted using a minimum of 10 adult zebrafish retinas or 6 mouse retinas at 2 days post-injury (dpi). Cells were lysed, and chromatin was sonicated to produce fragments ranging from 500 to 800 bp, following the method described by (Lindeman *et al*, 2009; Wang *et al*, 2023). The sonicated chromatin was divided into three equal portions: one served as the input control, another was immunoprecipitated using Rabbit polyclonal antibody against pSmad3 (Abcam, ab52903) or Rabbit polyclonal antibody against β-Catenin (Abcam, ab6302), and the third was immunoprecipitated with Rabbit IgG (Sigma Aldrich, I5006), which served as a negative control. The primers used for the ChIP assays are detailed in Table I.

### Luciferase assay

For luciferase assays, single-cell zebrafish embryos were injected with ∼1 nl of a solution containing 0.02 pg of *Renilla* luciferase mRNA (normalization), 5 pg of a *promoter*:GFP-*luciferase* vector (*malat1*, *mmp9*, *ascl1a*, *lin28a*, *her4.1*, *insm1a* promoter), and either *her4.1* mRNA, *her4.1* MO, *malat1* MO, SB431542 drug, or Tgf-β1 protein, as required. After 24 hours, embryos were divided into three groups (∼80 each) and lysed for dual-luciferase assays (Promega, E1910).

### Quantitative real-time RT–PCR, validation RT–PCR

Retinas were harvested after dark adaptation, and total RNA was extracted using TRIzol (Sigma). After DNase I (New England Biolabs, M0303L) treatment, double-stranded cDNA was synthesized from total RNA using a combination of random hexamers and oligo dT primers from cDNA synthesis kit (Thermo Fisher Scientific, K1622) or SuperScript-III (Thermo Fischer Scientific, 18080051). Quantitative real-time PCR was performed with at least six biological replicates, with each sample run in triplicate using the BIORAD SYBR mix on an Applied Biosystems real-time PCR system. Gene expression levels in control versus treated retinas were analyzed using the ΔΔCt method and normalized to *β-actin* or *l24* mRNA levels. The sequence of primers used for qPCR is provided in Table I.

### Western blotting and antibodies

Western blotting was performed using a single retina per experimental sample, lysed in Laemmli buffer. The lysate was size-fractionated on a 12% acrylamide gel under denaturing conditions and transferred to an Immuno-Blot polyvinylidene fluoride (PVDF) membrane (Biorad, cat. no. 162-0177). The membrane was then probed with specific primary antibodies and HRP-conjugated secondary antibodies for chemiluminescence detection using Clarity Western ECL (Bio-Rad, cat. no. 170-5061). Primary antibodies used in the study are rabbit polyclonal antibodies against human ASCL1/MASH1 (Abcam, cat. no. ab74065), Rabbit polyclonal antibody against LIN28a (Cell Signalling Technologies, cat. no. 3978), rabbit polyclonal antibody against Sox2 (cat. no. ab59776; Abcam), Rabbit polyclonal antibody against pSmad3 (Abcam, ab52903), Anti-YAP1 (phospho S127) antibody (EP1675Y), Rabbit polyclonal antibody against zebrafish Hdac1 (Harrison et al., 2011), Rabbit polyclonal antibody against H3K27ac (ab4729), Rabbit monoclonal antibody against Pten (138G6) (Cell Signaling Technologies, 9559). Gapdh was used as a loading control. HRP-conjugated secondary antibodies used were described previously (Kaur *et al*., 2018). For all Western blotting assays, 1–2 technical replicates were performed for each of the three biological replicates.

### BrdU/EdU labeling, tissue preparation, Immunofluorescence

BrdU labeling was done by immersing fish in 5 mM BrdU (Sigma) or injecting 10 μl of 20 mM BrdU solution intraperitoneally, 4 hours before euthanasia and retina dissection, unless otherwise mentioned. EdU labeling was performed via intraperitoneal injection of 10 mM EdU solution as described earlier (Mitra *et al*., 2018; Mitra *et al*, 2019). Fish were anesthetized, eyes dissected, lenses removed, fixed in 4% paraformaldehyde, cryopreserved, and sectioned (Fausett & Goldman, 2006). Immunofluorescence protocols and antibodies were as previously described (Kaur *et al*., 2018; Mitra *et al*., 2019; Ramachandran *et al*., 2010; Wan *et al*., 2012). Rat monoclonal antibody against BrdU (Abcam, catalog number ab6326); Mouse monoclonal antibody against human proliferating cell nuclear antigen, PCNA (Santa Cruz, catalog number sc-25280); Mouse polyclonal antibody against HuD (Santa Cruz, catalog number sc-48421); Goat polyclonal antibody against protein kinase C β1 (PKCβ1) (Santa Cruz, catalog number sc-209-G) were used. Secondary antibodies were conjugated to Alexa Fluor. EdU staining on cryosections was done with a Click-iT EdU Alexa Fluor 647 Imaging kit (Thermo Fisher Scientific, C10640). For all experiments, 3–6 retinae were used as biological replicates.

### RNA *in situ* hybridization

RNA *in situ* hybridization (ISH) was carried out on retinal sections using fluorescein- or digoxigenin-labeled complementary RNA probes (FL/DIG RNA Labeling Kit, Roche Diagnostics), following the protocol described in (Barthel & Raymond, 2000). Fluorescence ISH was performed according to the manufacturer’s instructions (Thermo Fisher Scientific, catalog numbers T20917, B40955, B40953).

### Strand-specific cDNA synthesis and ss-qPCR

Strand-specific cDNA was synthesized using a specifically recognized *talam1* primer with an attached tag, following the manufacturer’s protocol for gene-specific cDNA synthesis (Thermo Fisher Scientific, K1622). The 5′-tagged *talam1* cDNA was subsequently amplified, with the tag sequence as the forward primer and a *talam1*-specific reverse primer. For ss-qPCR, equal amounts of RNA were used to synthesize the first strand of *talam1* cDNA, while a separate aliquot was used to generate total cDNA using random primers and oligo-dT primers. qPCR was performed to analyze *talam1* expression, with normalization against β-actin using the corresponding total cDNA aliquot.

### FACS

GFP+ and GFP-cells were separated from injured *1016tuba1a:GFP* retinas at 4 dpi using a standardized protocol briefly outlined in (Ramachandran *et al*., 2011). 4 dpi injured retinas (40 each) from *1016tuba1a:GFP* were dissected in L15 media, followed by controlled hyaluronidase and trypsin treatment to prepare a single-cell suspension. The cells were sorted using a BD FACS Aria Fusion high-speed cell sorter. Total RNA was isolated from the sorted cells, and the levels of specific transcripts were quantified using qPCR.

### RNA Immunoprecipitation (RIP)

RIP assay was conducted using a minimum of 20 adult zebrafish 2 days post-injury (dpi) following the method described (Ren *et al*, 2020).

### Microscopy, cell counting, and statistical analysis

All slides after immunostaining or RNA *in situ* hybridization were mounted and examined using a Nikon Ni-E fluorescence microscope equipped with fluorescence optics and a Nikon A1 confocal imaging system. PCNA+ and BrdU+ cells were manually counted by observing fluorescence and ISH signals under a bright field. Statistical analysis was performed using a two-tailed unpaired t-test to compare two groups; for comparisons involving more than two groups, analysis of variance (ANOVA) was conducted, followed by a Dunnett test for post hoc analysis using GraphPad Prism. Error bars in all histograms represent standard deviation (s.d.).

## Acknowledgments

S.P. acknowledges her support from the IISER Mohali and CSIR for the Junior and Senior Research Fellowship. P.S. acknowledges postdoctoral fellowship support from DBT, RA (DBT-RA/2023/July/N/4351) and NPDF (PDF/2019/001148), DST India. P. Shukla acknowledges the CSIR Senior Research Fellowship. K. Y. acknowledges the UGC Senior Research Fellowship. O.M.D acknowledges support from IISER Mohali. R.R. also acknowledges research funding from DBT India (BT/PR53768/BMS/85/157/2024); (BT/PR36570/BRB/10/1976/2021) and support from IISER Mohali.

## Author contributions

R.R. conceived the study and designed experiments. S.P. performed most experiments, including conceptualization, methodology, data analysis, manuscript preparation, and review.

P.S. helped with mice sample acquisition and manuscript editing. M.S. helped in manuscript preparation and editing. K.Y helped in mice retina sample acquisition. O.M.D. contributed to the RIP protocol.

## EV-Figure legends

**Figure EV1: *malat1* is expressed in cells around proliferating cells, and is essential for zebrafish embryonic development and survival.**

**(A)** Confocal images after fluorescence *in-situ* hybridization of *malat1* at 4dpi, on retinal cross-section immunostained with BrdU, to mark proliferating cells at 4dpi. **(B)** Bright-field images of zebrafish embryos at 72 hours post-fertilization (hpf) demonstrating morphological defects following *malat1* knockdown using varying concentrations of morpholino. **(C)** Graph depicting the lethality rates of zebrafish embryos at 72 hpf upon injection with different concentrations of *malat1* morpholino. The data reveal a dose-dependent increase in lethality with increasing morpholino concentrations. Error bars represent standard deviation. *p < 0.005; n > 100 embryos analyzed per morpholino group. Each injection was repeated >5 times. Morpholino was injected into the yolk of one-cell stage embryos. Scale bar in **(A)**:10 µm and **(B)**:1mm.

**Figure EV2: *malat1* overexpression positively impacted MGPC proliferation, and rescued the effect of *malat1* MO.**

**(A)** The fluorescence image of a 36 hpf zebrafish embryo demonstrates reduced GFP expression upon injection with *malat1* MO and mRNA containing the *malat1* MO binding site (bs) upstream of *gfp*, compared to embryos injected with control MO and mRNA containing the *malat1* MO binding site (bs) upstream of *gfp***. (B)** Confocal images of retinal cross-sections at 4 dpi showed increased BrdU+ cells upon *malat1* RNA transfection, highlighting its pro-proliferative role. *malat1* knockdown reduces BrdU+ cells, an effect rescued by *malat1* overexpression. Data is presented as mean ± standard deviation (SD), *p < 0.05; n = 6 biological replicates. Quantification is shown in **(C)**. Scale bar: 1mm **(A)** and 10 µm **(B).**

**Figure EV3. *malat1* knockdown alters the expression of regeneration-associated genes and affects chromatin status.**

**(A)** Luciferase assay in zebrafish embryos (24hpf) shows dysregulation in *ascl1a, lin28a, mmp9, her4.1,* and *insm1a* promoter activity on *malat1* knockdown. Data is presented as mean ± standard deviation (SD), *p < 0.04; n = 6 biological replicates. **(B)** Western blot analysis shows changes in the expression of regeneration-associated proteins, epigenetic modifiers, signaling molecules, and epigenetic marks on chromatin following *malat1* knockdown at 2 dpi. These findings suggest that *malat1* impacts a wide range of proteins and modulates the chromatin status of cells to mediate its effects. **(C)** A schematic depicting the arrangement of the *malat1* gene and its neighboring genes on linear DNA in humans, mice, and zebrafish.

**Figure EV4: *malat1* positively affects *egr1* expression, a pro-proliferative factor during zebrafish retina regeneration.**

**(A)** qPCR analysis of *egr1* expression levels at various time points post-retinal injury, showing early induction of *egr1* after injury. Data is presented as mean ± standard deviation (SD), *p < 0.005; n = 6 biological replicates. **(B)** Confocal images of retinal cross-sections from the *tuba1a1016:gfp* transgenic line at 4 dpi, showing a decline in GFP-positive proliferating cells upon *egr1* knockdown in a concentration-dependent manner, quantified in **(C).** Data is presented as mean ± standard deviation (SD), *p < 0.0005; n = 6 biological replicates. **(D)** Bright-field images of *malat1 in-situ* hybridization following *egr1* knockdown and overexpression, respectively, indicating that *egr1* positively regulates *malat1*, suggesting a positive feedback loop, quantified in **(E)**. Data is presented as mean ± standard deviation (SD), *p < 0.0005; n = 6 biological replicates. Scale bars represent 10 µm **(B, D)**.

**Fig EV5: Egr1 can induce MGPC proliferation but requires sufficient induction of RAGs for effective regeneration to happen.**

**(A)** Confocal images of retinal cross-sections showing PCNA immunostaining in retinas where *egr1* was overexpressed without injury, indicating MGPC proliferation occurs even without injury, if *egr1* is overexpressed. Quantification is shown in **(B)**. Data is presented as mean ± standard deviation, *p < 0.0003; n = 6 biological replicates. **(C)** qPCR analysis of regeneration-associated genes (RAGs) at 2 days post-transfection (dpt) with *egr1* overexpression in uninjured retinae. No significant induction of RAGs was observed (ns indicates non-significant). Data is presented as mean ± standard deviation, *p < 0.05; n = 6 biological replicates. **(D)** Schematic representation of the experimental regime for evaluating the viability of proliferated MGPCs (cells labeled with BrdU at 3/4/5 dpt) after *egr1* overexpression without injury. **(E)** Confocal images of retinal sections show that while *egr1* overexpression induced MGPC proliferation at 6 dpt, these proliferated cells were not viable by 30 days post-transfection (30dpt), as no BrdU+ cells were observed at 30 dpt. **(F)** Quantification of MGPC proliferation and viability at 6 dpt and 30 dpt respectively, on *egr1* overexpression without injury. Data is presented as mean ± standard deviation, *p < 0.005; n = 6 biological replicates. Scale bars represent 10 µm (**A, E)**.

**Figure EV6: Wnt signaling regulates *egr1* during retina regeneration.**

**(A)** Confocal images illustrating that the effect of Wnt signaling inhibition (using XAV939) on reducing proliferation is rescued by overexpressing *egr1*, quantified in **(B)**. Data is presented as mean ± standard deviation, *p < 0.0003; n = 6 biological replicates. Scale bars represent 10 µm. **(C)** qPCR analysis shows that *egr1* gets induced by stabilizing β-catenin (by SB216763 administration), with or without injury. Conversely, *egr1* levels decline upon Wnt signaling inhibition (XAV939 administration), demonstrating a critical role for Wnt signaling in modulating *egr1* expression. *p< 0.05; n=6 biological replicates. Error bars represent standard deviation.

**Figure EV7: Notch signaling and Wnt signaling regulate *malat1* expression.**

**Neighboring cells express *malat1* and *dlld*, while proliferating cells express *her4.1* (A)** Luciferase assay in zebrafish embryos (24hpf) supports that *her4.1* knockdown upregulates *malat1* expression, and *her4.1* overexpression suppresses *malat1* expression. **(B)** Double Fluorescence *in-situ* hybridization (FISH) shows *malat1* expressing cells have more *dlld* expression and **(C)** *malat1* expressing cells and *her4.1* expressing cells lie adjacent to each other. **(D)** qPCR analysis in sorted cells, from 4dpi injured *tuba1016:gfp* transgenic retina, where GFP+ve fraction contains proliferating cells, and GFP-ve fraction contains non-proliferating cells, show *her4.1* enrichment in GFP +ve (actively proliferating) and *dlld* enrichment in GFP-ve (neighboring) fraction. Data is shown as mean values ± SD, *p < 0.05; n = 6 biological replicates. **(E)** qPCR analysis from the sorted cells reveal enrichment of *egr1* transcript in GFP+ fraction (actively proliferating cells). Data is shown as mean values ± SD, *p < 0.05; n = 6 biological replicates.

**Figure EV8: Differential regulation of *malat1* by TGF-β signaling in zebrafish and mice is exerted through direct binding of pSmad3 on regulatory elements of *talam1* promoter. TGF-β signaling regulates *egr1* expression in both zebrafish and mice like its regulation of *malat1*.**

**(A)** Luciferase assay in zebrafish embryos (24hpf) supports that SB431542 (TGF-β inhibitor) and TGF-β1 overexpression upregulates and downregulates *malat1* expression, respectively. Data is shown as mean values ± SD, *p < 0.003; n = 6 biological replicates. **(B)** Orientation of *malat1* and *talam1* genes. **(C)** Diagrammatic representation of TIE elements on the zebrafish *talam1* promoter. **(D)** PCR analysis after pSmad3 ChIP in 2 dpi zebrafish retinae shows pSmad3 occupancy on TIE elements of the *talam1* promoter. **(E)** Diagram showing 5GC and TIE elements on the mouse *Malat1* promoter. **(F)** PCR after PSmad3 ChIP in mouse retinae shows PSmad3 occupancy on 5GC and TIE elements of the *Malat1* promoter. **(G)** qPCR analysis showing downregulation of *egr1* in zebrafish, *p < 0.03; n = 6 biological replicates and **(H)** upregulation of *Egr1* in mice upon TGF-β1 overexpression, *p < 0.05; n = 6 biological replicates. Data is shown as mean values ± SD. **(I)** ssRT-PCR showing the presence of *talam1* transcript in zebrafish embryos, with genomic DNA as a positive control. **(J)** Fluorescent image of a 36 hpf *talam1* zebrafish reporter line showing GFP expression along the CNS, indicating *talam1* promoter activity in embryos. **(K)** transgenic *malat1* reporter line embryo, showing similar expression of GFP along CNS, albeit stronger than the *talam1* reporter line. Scale bar: 1mm **(J)** and **(K).**

**Figure EV9: A schematic representation of the gene regulatory network elucidated in this study.**

